# Infection-dependent antibiotic response phenotypes emerge during chronic airway adaptation of *Pseudomonas aeruginosa*

**DOI:** 10.64898/2026.11.11.765915

**Authors:** Pablo Laborda, Esmeralda Cebrián-Sastre, Ruggero La Rosa, Søren Molin, Helle Krogh Johansen

## Abstract

**Objectives:** Chronic *Pseudomonas aeruginosa* airway infections can persist for years despite intensive antibiotic therapy guided by standard antimicrobial susceptibility testing. While this technique captures the emergence of resistance, it does not reproduce many features of the infection environment, where adaptation acquired during chronic infection may alter bacterial responses to antibiotics in ways that are not fully reflected by conventional susceptibility testing. Here we investigated antibiotic responses of longitudinal clinical isolates under standard laboratory conditions and in an airway epithelial infection model.

**Methods:** Six longitudinal pairs of *P. aeruginosa* isolates from people with cystic fibrosis were analysed. Changes in bacterial growth following exposure to tobramycin, ceftazidime and ciprofloxacin were quantified under standard laboratory conditions and compared with changes in bacterial burden after treatment measured in differentiated airway epithelial cultures maintained under air-liquid interface (ALI) conditions that reproduce key features of the airway infection environment. Representative phenotypes were visualized by confocal microscopy and interpreted in the context of available whole-genome sequencing data.

**Results:** Major resistance phenotypes identified by conventional susceptibility testing were largely reproduced in the ALI model. However, concordance between phenotypes observed under standard laboratory and ALI conditions differed between antibiotics. Several isolates classified as susceptible according to conventional susceptibility testing displayed substantial survival in the ALI model, and these discordant phenotypes were observed predominantly among those isolates recovered after prolonged airway persistence. Genomic analyses identified mutations associated with both classical resistance mechanisms and chronic adaptation.

**Conclusions:** Phenotypes observed under conventional laboratory conditions do not uniformly predict those observed in the ALI airway conditions. Discordant phenotypes were most apparent among chronically adapted isolates and varied between antibiotics, suggesting that adaptation acquired during chronic infection can alter antibiotic responses in ways that are not fully reflected by conventional susceptibility testing.

## Introduction

*Pseudomonas aeruginosa* is a major cause of chronic airway infections in individuals with underlying lung diseases such as cystic fibrosis (CF) [1]. Once established, infections can persist for years despite intensive and repeated antibiotic treatment, creating selective pressures that drive bacterial adaptation within the host. As a result, genetically and phenotypically distinct variants emerge throughout the course of infection, shaping long-term persistence and treatment outcomes [2–4].

Resistance acquired during chronic infection is typically assessed through antimicrobial susceptibility testing based on minimum inhibitory concentration (MIC) measurements [5]. Although MIC testing reliably detects many clinically important resistance mechanisms, increasing evidence suggests that antibiotic activity can differ substantially under infection-relevant conditions compared with standard laboratory media [6,7]. Consequently, in addition to conventional resistance, adaptation during chronic infection may alter how bacteria respond to antibiotics, generating phenotypes that become apparent only under infection-relevant conditions. Consistent with this idea, reconstruction of adaptive mutations identified in *P. aeruginosa* clinical isolates has revealed altered antibiotic responses through mechanisms linked to infection-specific physiology, tissue colonization, and host-pathogen interactions [8–10].

To investigate this possibility, we compared early and late *P. aeruginosa* clinical isolates longitudinally collected from chronically infected people with CF. Changes in bacterial growth and survival following antibiotic exposure were assessed under conventional laboratory conditions and in a differentiated human airway epithelial air-liquid interface (ALI) infection model that reproduces key features of the respiratory environment [11]. By directly comparing these approaches, we sought to determine whether adaptation during chronic infection is associated with infection-dependent antibiotic response phenotypes and how these can vary across isolates and antibiotic treatments.

## Methods

### Bacterial strains and culture conditions

The *P. aeruginosa* clinical isolates used in this study have been previously described and whole genome sequenced [2], except for DK08 (strains 511.1 and 512), which was sequenced subsequently. Mutations in strain 511.1 were identified by comparative analysis against the genome assembly of strain 512 (ENA accession SAMEA3176788) using breseq v0.40.1 [12].

The use of stored bacterial isolates was approved by the Ethics Committee of the Capital Region of Denmark (Region Hovedstaden; approval number H-21078844).

Bacterial strains were routinely cultured in Lysogeny Broth (LB) at 37°C under aerobic conditions.

### Antibiotic susceptibility testing

Minimum inhibitory concentrations (MICs) of ciprofloxacin, ceftazidime and tobramycin were determined by broth microdilution in Mueller-Hinton (MH) broth according to EUCAST recommendations [13]. Optical density (OD) at 600 nm was measured at the end of the assay using a BioTek ELx808 microplate reader (BioTek Instruments).

To enable direct comparison of bacterial responses to antibiotic treatment in MH broth and the ALI infection model, bacterial growth was additionally quantified by CFU enumeration in MH broth containing subinhibitory antibiotic concentrations after 24 h incubation. Ciprofloxacin, tobramycin and ceftazidime were tested at 0.2, 0.5 and 1 μg/mL, respectively. These concentrations permitted growth of all isolates under a common treatment condition. All experiments were performed in three independent biological replicates.

### Generation of GFP-expressing strains

Strains were chromosomally tagged with GFP by Tn7-mediated insertion using plasmid pUC18T-mini-Tn7T-Gm-Tp-gfp as previously described [8,14,15]. Successful tagging was confirmed by fluorescence microscopy.

### Air-liquid interface airway epithelial cultures

ALI cultures were established using the human bronchial epithelial cell line BCi-NS1.1 [16] following a previously described protocol [8,9,11,15,17]. Briefly, 1 × 10⁵ cells were seeded onto collagen-coated 1-µm pore polyester membrane inserts (Falcon) and differentiated at an air-liquid interface for 28 days in PneumaCult™-ALI medium (STEMCELL Technologies) at 37°C and 5% CO₂. Airway epithelial differentiation and barrier formation were monitored by transepithelial electrical resistance (TEER) measurements using an STX2 electrode (World Precision Instruments). To prevent mucus accumulation, the apical surface was washed weekly with PBS from day 15.

### ALI infection model and antibiotic treatment

Overnight bacterial cultures were diluted to an OD₆₀₀ of 0.01 and grown to mid-exponential phase. Cells were washed and adjusted to 1 × 10⁵ CFU/mL in PBS. Differentiated ALI cultures were infected apically with approximately 1 × 10³ CFU and incubated for 7 h at 37°C and 5% CO₂.

Antibiotics were subsequently added to the basolateral compartment at 4 μg/mL tobramycin, 8 μg/mL ceftazidime or 0.5 μg/mL ciprofloxacin, and cultures were incubated for an additional 14 h. Bacterial burden before and after treatment was measured by CFU quantification from apical washes, basolateral medium and epithelial-associated bacteria recovered by scraping the epithelial layer. Total bacterial burden was calculated as the sum of CFUs recovered from all compartments.

Antibiotic response was defined as the fold change in total bacterial number after treatment relative to the number measured immediately before antibiotic exposure. To assess adaptation during chronic infection, treatment responses of late isolates were normalized to those of their corresponding early isolates. All experiments were performed in three independent biological replicates.

### Confocal microscopy

ALI cultures were infected and treated as described above, fixed with 4% paraformaldehyde, stained with Phalloidin-AF555 and TO-PRO-3, and imaged using a Leica Stellaris 8 confocal microscope as previously described [8,9,11,15,17]. GFP-expressing bacteria were visualized together with epithelial actin and nuclei staining.

### Statistical analysis

Statistical analyses were performed using GraphPad Prism v10.3.1 (GraphPad Software). Antibiotic treatment data measured in MH broth and the ALI model were log_10_-transformed prior to analysis. For each clone type, differences between early and late isolates across antibiotic treatments were evaluated using two-way ANOVA followed by Šídák’s multiple-comparisons test. Associations between phenotypes measured under standard laboratory conditions and in the ALI infection model were evaluated using Pearson correlation analysis. Statistical significance was defined as *P* < 0.05. Figures were generated using Adobe Illustrator v29.7 (Adobe).

## Results

### Antibiotic treatment phenotypes under standard laboratory and infection-relevant conditions show variable discordance across antibiotics

To investigate whether adaptation acquired during chronic infection is associated with phenotypes that are not fully reflected by conventional susceptibility testing, six pairs of longitudinal *P. aeruginosa* isolates recovered between 2004-2012 from people with CF were selected. Each pair comprised an early isolate recovered shortly after detection of the infecting clone type and a late isolate obtained from the same lineage after 2.6–7.0 years of persistence in the airways (Table 1). The selected lineages represented diverse evolutionary trajectories of resistance acquisition to tobramycin, ceftazidime and ciprofloxacin, three frequently used anti-pseudomonal antibiotics, according to conventional susceptibility testing (Table 1; Supplementary Figure S1).

**Table 1.**
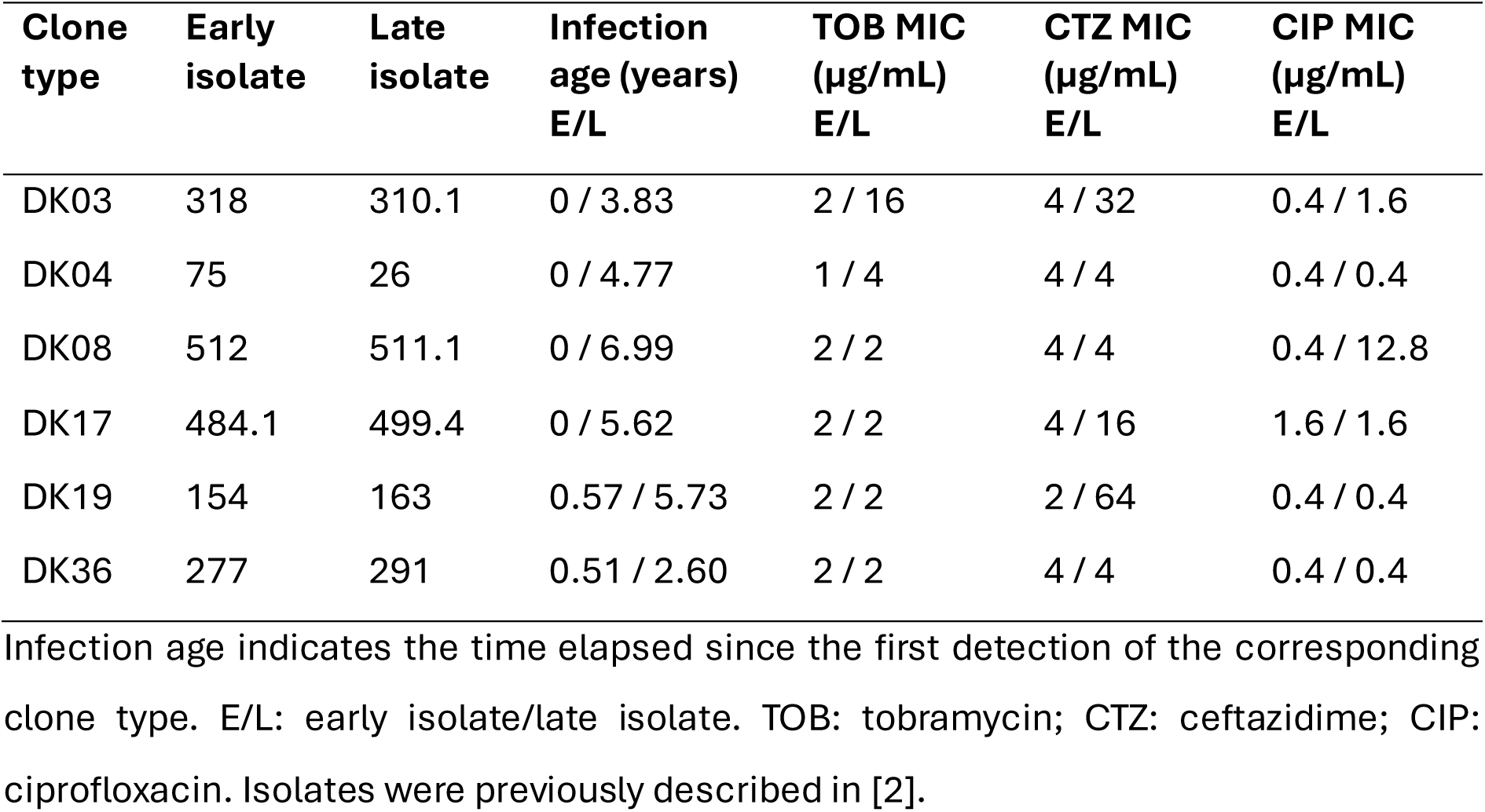
Characteristics and MIC profiles of longitudinal *Pseudomonas aeruginosa* clinical isolate pairs included in this study.

| **Clone type** | **Early isolate** | **Late isolate** | **Infection age (years)**<br>**E/L** | **TOB MIC (µg/mL)**<br>**E/L** | **CTZ MIC (µg/mL)**<br>**E/L** | **CIP MIC (µg/mL)**<br>**E/L** |
|---|---|---|---|---|---|---|
| DK03 | 318 | 310.1 | 0 / 3.83 | 2 / 16 | 4 / 32 | 0.4 / 1.6 |
| DK04 | 75 | 26 | 0 / 4.77 | 1 / 4 | 4 / 4 | 0.4 / 0.4 |
| DK08 | 512 | 511.1 | 0 / 6.99 | 2 / 2 | 4 / 4 | 0.4 / 12.8 |
| DK17 | 484.1 | 499.4 | 0 / 5.62 | 2 / 2 | 4 / 16 | 1.6 / 1.6 |
| DK19 | 154 | 163 | 0.57 / 5.73 | 2 / 2 | 2 / 64 | 0.4 / 0.4 |
| DK36 | 277 | 291 | 0.51 / 2.60 | 2 / 2 | 4 / 4 | 0.4 / 0.4 |

Infection age indicates the time elapsed since the first detection of the corresponding clone type. E/L: early isolate/late isolate. TOB: tobramycin; CTZ: ceftazidime; CIP: ciprofloxacin. Isolates were previously described in [2].

Bacterial growth under antibiotic exposure was quantified in MH broth and also in differentiated airway epithelial cultures using an ALI infection model that reproduces key features of the airway environment [11] (Figure 1). For each assay, identical antibiotic concentrations were applied to all isolates to enable comparison across lineages. Antibiotic concentrations were selected to impose measurable growth inhibition without completely eliminating the infecting population. In both systems, bacterial growth and survival after antibiotic exposure were quantified by CFU enumeration.

**Figure 1.**
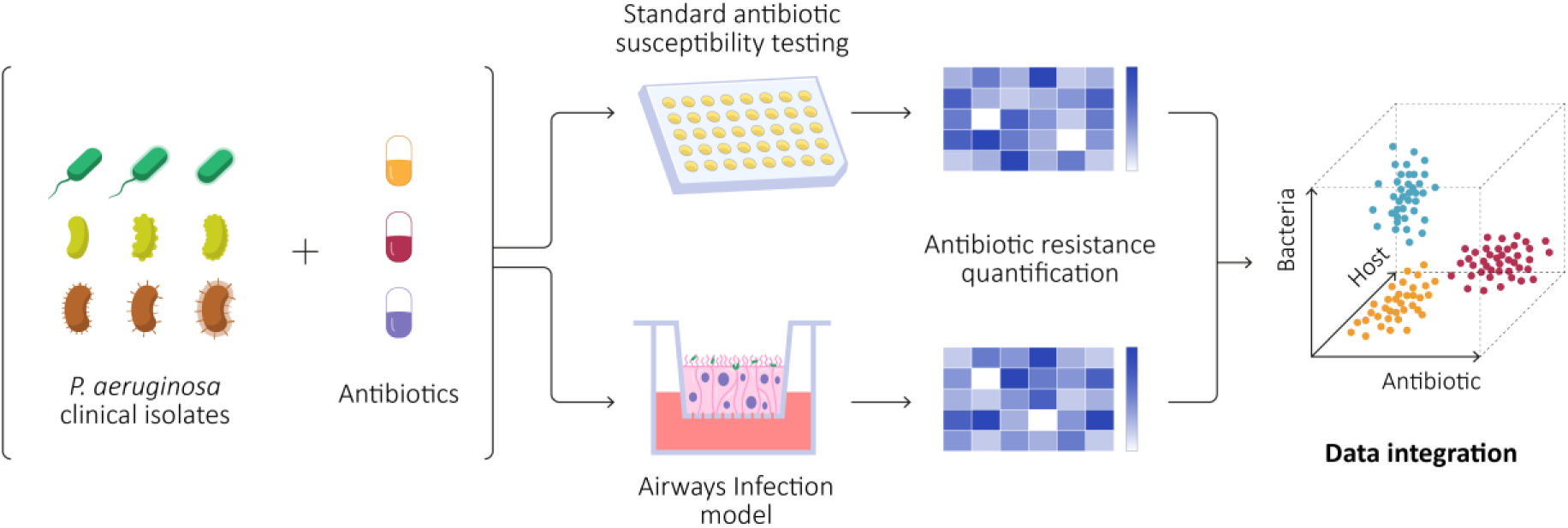
Experimental workflow for the analysis of infection-dependent antibiotic responses. Longitudinal *Pseudomonas aeruginosa* clinical isolates representing distinct stages of chronic infection were exposed to tobramycin, ceftazidime and ciprofloxacin. Antibiotic responses were evaluated under standard laboratory conditions and in differentiated airway epithelial cultures maintained under air-liquid interface conditions. Integration of these datasets enabled comparison of antibiotic responses across bacterial lineages, antimicrobial treatments and environmental conditions, revealing infection-dependent antibiotic response phenotypes associated with chronic adaptation.

To assess the relationship between both experimental conditions, the measured phenotypes were compared for each isolate and antibiotic combination (Figure 2). For tobramycin, phenotypes observed in MH broth and in the ALI model showed a moderate significant positive correlation (Pearson r = 0.83, P = 0.0008), although several isolates displayed discordant phenotypes between environments. In contrast, ceftazidime and ciprofloxacin showed weaker, non-significant associations (ceftazidime: Pearson r = 0.48, P = 0.1186; ciprofloxacin: Pearson r = 0.51, P = 0.0868) and revealed multiple discordant phenotypes. While isolates classified as resistant according to conventional susceptibility testing displayed phenotypes consistent with reduced antibiotic efficacy in both systems, several isolates classified as susceptible nevertheless exhibited substantial survival in the ALI model.

**Figure 2.**
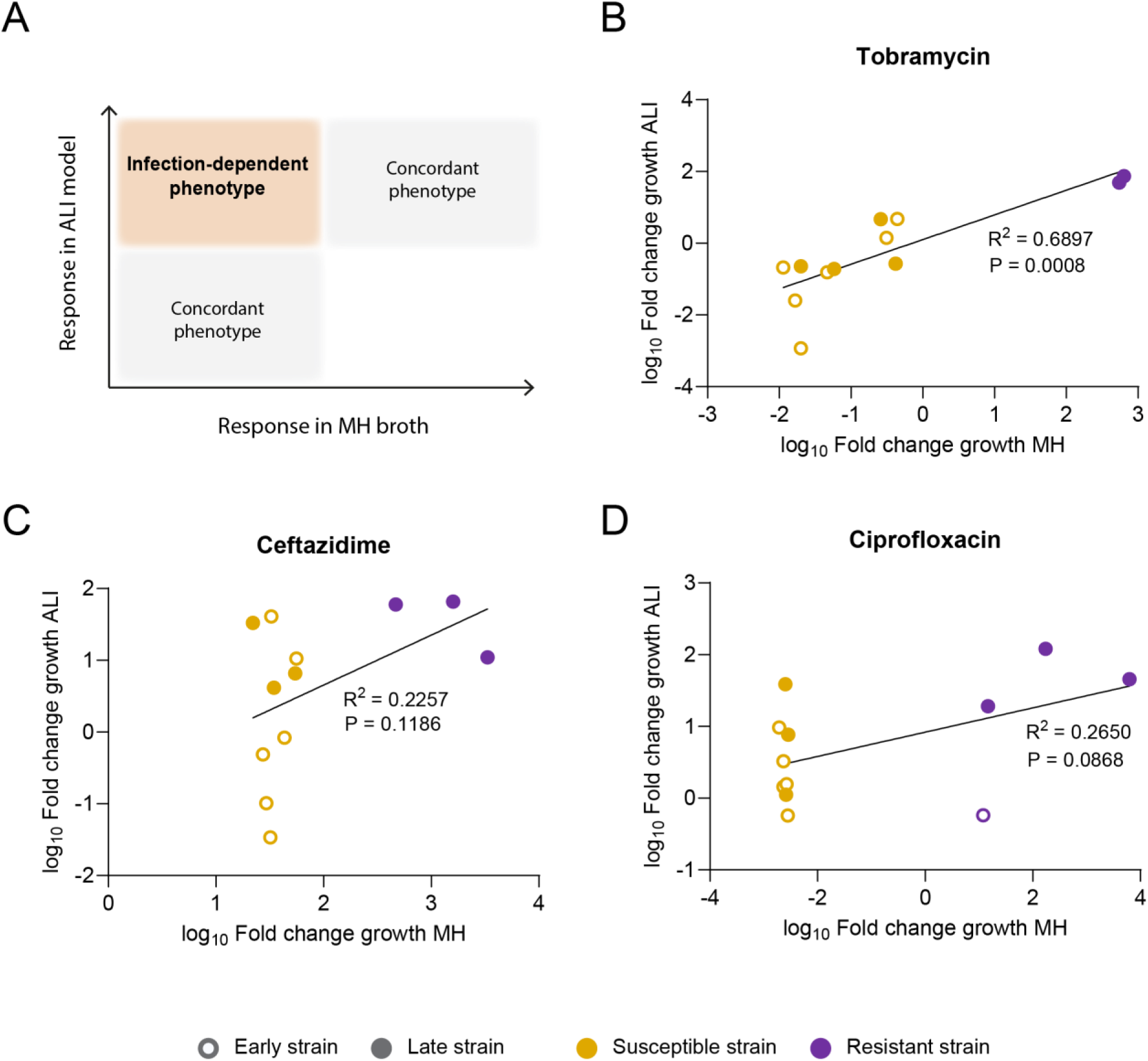
Correlation between phenotypes observed under standard laboratory and infection-relevant conditions. **(A)** Schematic representation of concordant and discordant antibiotic response phenotypes across experimental environments. **(B-D)** Antibiotic response phenotypes of longitudinal *Pseudomonas aeruginosa* clinical isolates in Mueller-Hinton (MH) broth and in differentiated airway epithelial cultures maintained under air-liquid interface (ALI) conditions. Each point represents an individual isolate. The x-axis shows the log₁₀-transformed fold change in bacterial growth measured in MH broth containing subinhibitory concentrations of tobramycin **(B)**, ceftazidime **(C)**, or ciprofloxacin **(D)**, while the y-axis shows the corresponding log₁₀-transformed fold change in bacterial burden measured in the ALI infection model following antibiotic treatment. Isolates classified as susceptible or resistant according to EUCAST clinical breakpoints are shown in yellow and purple, respectively. Early isolates are represented by open circles and late isolates by filled circles. Linear regression lines, coefficients of determination (R²) and P values are shown. Concordance between phenotypes measured under standard laboratory conditions and in the ALI model differed between antibiotics, with several isolates displaying discordant phenotypes across environments.

Collectively, these observations indicate that phenotypes observed under standard laboratory conditions do not uniformly predict those observed in an airway epithelial infection model and suggest that the extent of this discordance differs between antibiotics.

### Infection-dependent antibiotic response phenotypes emerge during chronic infection

To further investigate the extent to which adaptation during chronic infection contributed to the discordant phenotypes identified above, antibiotic treatment phenotypes of late isolates were compared with those of their corresponding early isolates under both standard laboratory and infection-relevant conditions (Figure 3).

**Figure 3.**
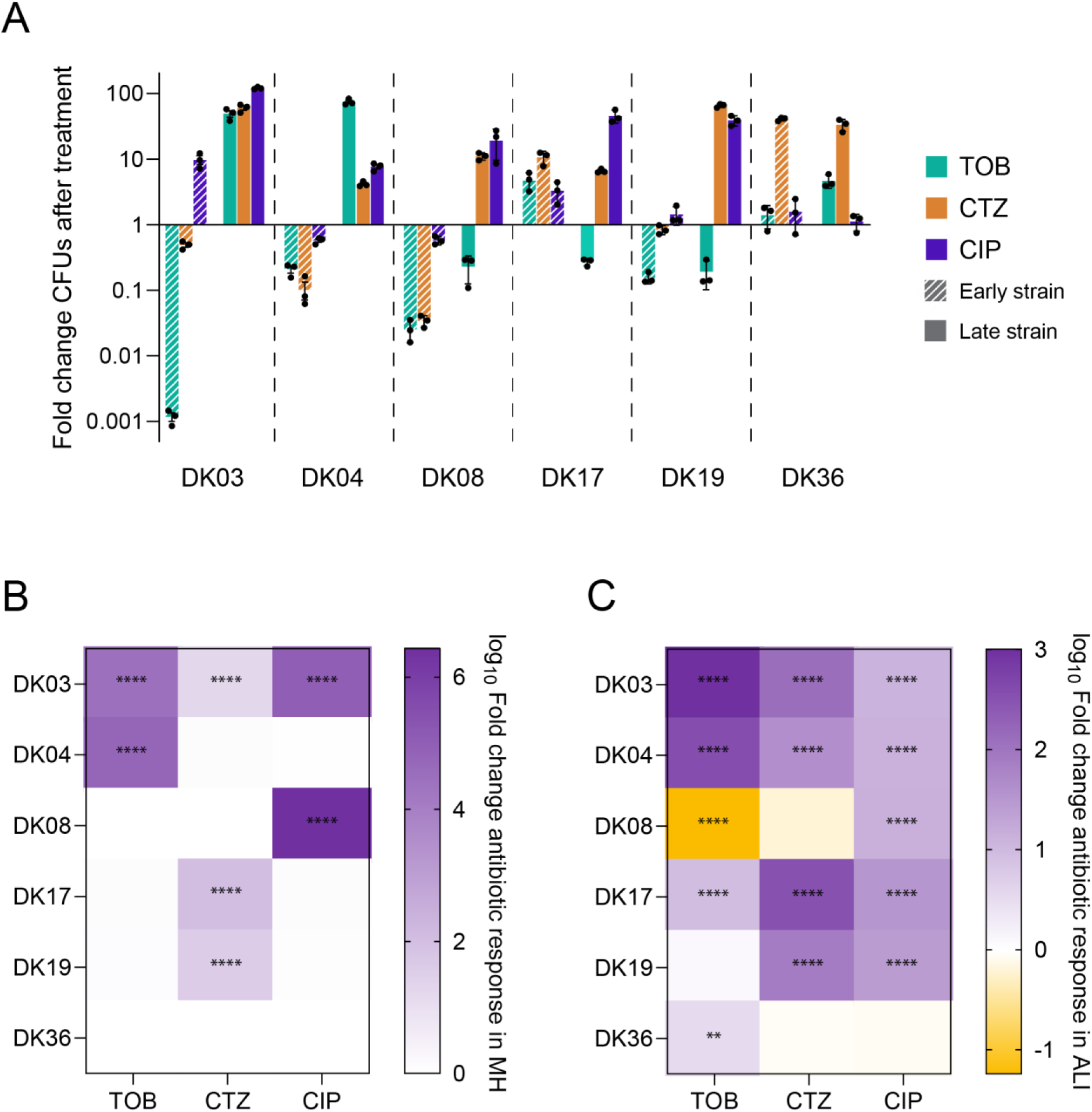
Comparison of antibiotic response changes acquired during chronic infection under standard laboratory and infection-relevant conditions. **(A)** Antibiotic responses in the air-liquid interface (ALI) infection model, expressed as the fold change in bacterial burden after treatment relative to the burden immediately before antibiotic exposure. ALI cultures were infected with the indicated isolates and bacterial burden was determined by CFU enumeration before and after antibiotic treatment. Data are shown for early (striped bars) and late (solid bars) isolates from each clone type. Bars represent mean values and error bars indicate the standard error of the mean (SEM). Individual data points represent biological replicates. Colours indicate antibiotic treatments: tobramycin (TOB, green), ceftazidime (CTZ, orange) and ciprofloxacin (CIP, purple). Values greater than 1 indicate an increase in bacterial burden relative to pre-treatment levels, whereas values below 1 indicate a reduction in bacterial burden following antibiotic exposure. Raw data are provided in Supplementary Figure S2. **(B)** Heatmap showing changes in antibiotic responses measured in Mueller Hinton (MH) broth. Values represent the log10-transformed ratio of antibiotic responses in late isolates relative to their corresponding early isolates following exposure to subinhibitory concentrations of tobramycin, ceftazidime, or ciprofloxacin in MH broth. Antibiotic responses were quantified as the fold change in CFUs after relative to before antibiotic treatment. Positive values indicate enhanced growth of the late isolate relative to the corresponding early isolate under antibiotic treatment. **(C)** Heatmap showing changes in antibiotic responses measured in the ALI infection model. Values represent the log10-transformed ratio of the treatment response of late isolates relative to their corresponding early isolates, calculated from the data shown in panel A. Positive values indicate enhanced survival of the late isolate relative to the corresponding early isolate following antibiotic treatment. Statistical significance is indicated by asterisks within the corresponding heatmap cells and was determined from the raw data underlying each assay. Data were log10-transformed prior to analysis and compared using two-way ANOVA followed by Šídák’s multiple-comparisons test. P < 0.01 (**), P < 0.0001 (****).

Conventional susceptibility testing revealed diverse resistance trajectories, including increased ceftazidime resistance in DK19 and DK17, multidrug resistance in DK03, increased tobramycin resistance in DK04, increased ciprofloxacin resistance in DK08, or no major resistance changes in DK36 (Table 1; Supplementary Figure S1). When changes in bacterial burden following antibiotic treatment were quantified in the ALI model, the major resistance phenotypes identified by conventional testing were largely recapitulated. However, comparison of the two conditions also identified phenotypes that were only apparent in the ALI model. Enhanced survival after antibiotic treatment was detected for several late isolates despite showing limited changes in MH broth, particularly during ciprofloxacin treatment of DK19 and DK17 and during ceftazidime and ciprofloxacin treatment of DK04 (two-way ANOVA with Šídák’s multiple-comparisons test, *P* < 0.0001) (Figure 3B, C).

Two isolate pairs displaying marked discordance between standard laboratory and ALI phenotypes were selected for confocal microscopy: DK17 treated with ciprofloxacin and DK04 treated with ceftazidime. In both cases, late isolates formed larger and/or more abundant bacterial aggregates following antibiotic treatment than their corresponding early isolates (Figure 4). These observations were consistent with CFU quantification and provided independent visual confirmation of the enhanced survival phenotypes identified in the ALI infection model.

**Figure 4.**
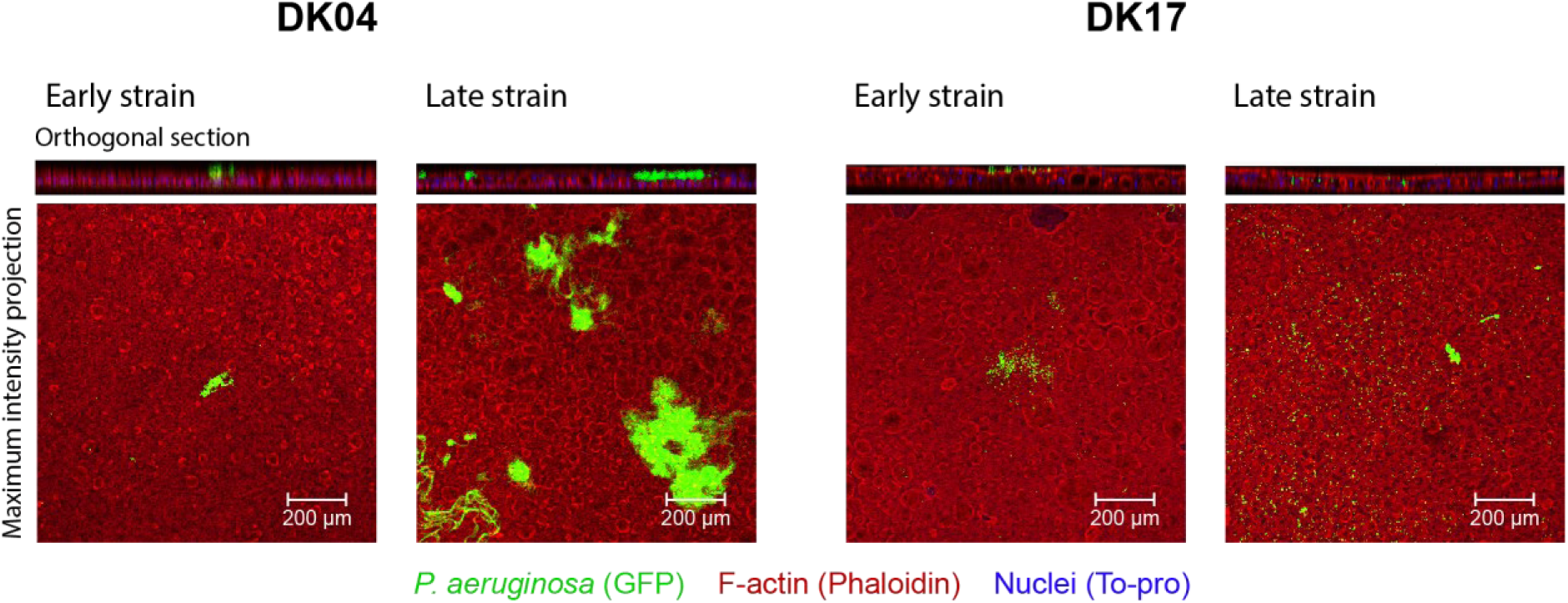
Representative confocal microscopy images of longitudinal isolate pairs following antibiotic treatment in the airway infection model. Differentiated airway epithelial cultures were infected with GFP-expressing *Pseudomonas aeruginosa* isolates and treated with ceftazidime (DK04) or ciprofloxacin (DK17). Representative images of early and late isolates are shown. Bacteria are shown in green (GFP), epithelial actin in red (phalloidin), and host cell nuclei in blue (TO-PRO-3). Nuclei signal was omitted from maximum intensity projections for image clarity. Larger and/or more abundant bacterial aggregates were observed for late isolates compared with their corresponding early isolates, consistent with the enhanced survival phenotypes identified by CFU quantification in the air-liquid interface infection model. Scale bar: 200 μm. Images are representative of three independent experiments.

These findings indicate that infection-dependent antibiotic response phenotypes emerge during chronic infection and are incompletely reflected by conventional laboratory susceptibility measurements. Notably, these phenotypes were observed predominantly in late isolates, suggesting that prolonged adaptation to the host environment increases the likelihood of discordant phenotypes across experimental conditions.

### Genetic changes accumulated during chronic infection

Whole-genome comparison of longitudinal isolate pairs identified numerous genetic changes accumulated during chronic infection (Supplementary Table S1). Several mutations previously associated with antibiotic resistance were detected in lineages displaying corresponding standard susceptibility measurements changes. For example, DK19 carried mutations in *mexB* and *mpl*, both previously linked to β-lactam resistance [18–20]. Similarly, DK08 acquired a mutation in *nfxB*, a well-characterized regulator associated with fluoroquinolone resistance [19], while a mutation in *fusA1* was identified in the tobramycin-resistant late DK03 isolate [21]. In contrast, DK19 also acquired a mutation in *gyrB*, encoding the target of ciprofloxacin [22], despite showing no increase in ciprofloxacin MIC while exhibiting enhanced survival in the ALI model.

Several lineages also accumulated mutations previously associated with chronic airway adaptation, including alterations in *lasR*, *retS*, *gacS* and other global regulatory genes [2]. Notably, *mexZ* was mutated in four lineages (DK19, DK17, DK03 and DK04), three of which displayed enhanced survival in the ALI model without corresponding MIC changes.

Together, these findings suggest that the phenotypes observed in the ALI model are associated with both classical antibiotic resistance determinants and mutations linked to long-term adaptation to the airway environment.

## Discussion

Chronic airway infections caused by *P. aeruginosa* remain difficult to eradicate despite intensive antibiotic therapy [1,3,10]. Clinical treatment decisions are largely guided by conventional antibiotic susceptibility testing [5], yet persistent infections can occur even when infecting isolates remain classified as susceptible [10]. Previous studies have reported that only a small proportion (2-10%) of *P. aeruginosa* clinical isolates are classified as resistant when recovered from chronically infected patients [10], suggesting that factors contributing to antibiotic treatment failure are not always captured by conventional MIC testing alone [8,9,23]. In the present study, we compared phenotypes observed under standard laboratory conditions with those observed in an airway epithelial infection model. While major resistance phenotypes identified by conventional susceptibility testing were largely reproduced in the ALI model, additional phenotypes emerged under infection-relevant conditions.

An important observation was that the degree of concordance differed between antibiotics. While tobramycin treatment phenotypes observed under standard laboratory conditions showed a significant correlation with those observed in the ALI model, ceftazidime and ciprofloxacin revealed weaker associations and multiple discordant phenotypes. Notably, several isolates classified as susceptible according to conventional susceptibility testing nevertheless displayed substantial survival in the ALI model. Although this proof-of-concept study included a limited number of longitudinal lineages, precluding broader conclusions regarding antibiotic-specific patterns, these findings suggest that the relationship between conventional susceptibility measurements and infection-relevant antibiotic responses may depend on the antimicrobial context [8,9].

Discordant phenotypes were observed predominantly among isolates recovered after prolonged persistence in the airways. This suggests that adaptation acquired during chronic infection increases the likelihood that antibiotic responses become dependent on environmental context. Importantly, conventional resistance phenotypes remained detectable under both experimental conditions, indicating that classical resistance mechanisms remain major determinants of antibiotic responses across environments. However, the additional phenotypes revealed by the ALI model demonstrate that chronic adaptation can influence bacterial responses to antibiotic treatment in ways incompletely reflected by standard laboratory measurements.

These findings support the emerging view that persistence during chronic airway infection extends beyond conventional antibiotic resistance and is shaped by host-pathogen interactions [6,8–10]. Adapted *P. aeruginosa* populations can exploit multiple strategies to survive during infection, including metabolic adaptation, altered regulatory networks, tissue-associated growth, biofilm-associated growth and other persistence mechanisms that are largely absent from standard susceptibility assays. Notably, the phenotypes identified here emerged under conditions that capture early host-pathogen interactions, suggesting that adaptation to the host environment can influence antibiotic survival from the earliest stages of infection. As a result, antibiotic treatment failure may arise not only through acquisition of classical resistance determinants but also through adaptations that become relevant only within the host environment, highlighting the value of infection-relevant models for studying bacterial persistence [24]. Consistent with this notion, all six lineages studied here remained established in the same people with CF until at least 2020, up to more than a decade after the isolates analyzed here were collected, despite largely retaining susceptibility to routinely monitored antibiotics (Table 2). Notably, these patients were managed according to intensive Danish CF treatment protocols involving repeated antibiotic intervention [2,25]. Although not investigated in this study, this observation illustrates that prolonged persistence can occur without extensive conventional resistance phenotypes.

**Table 2.**
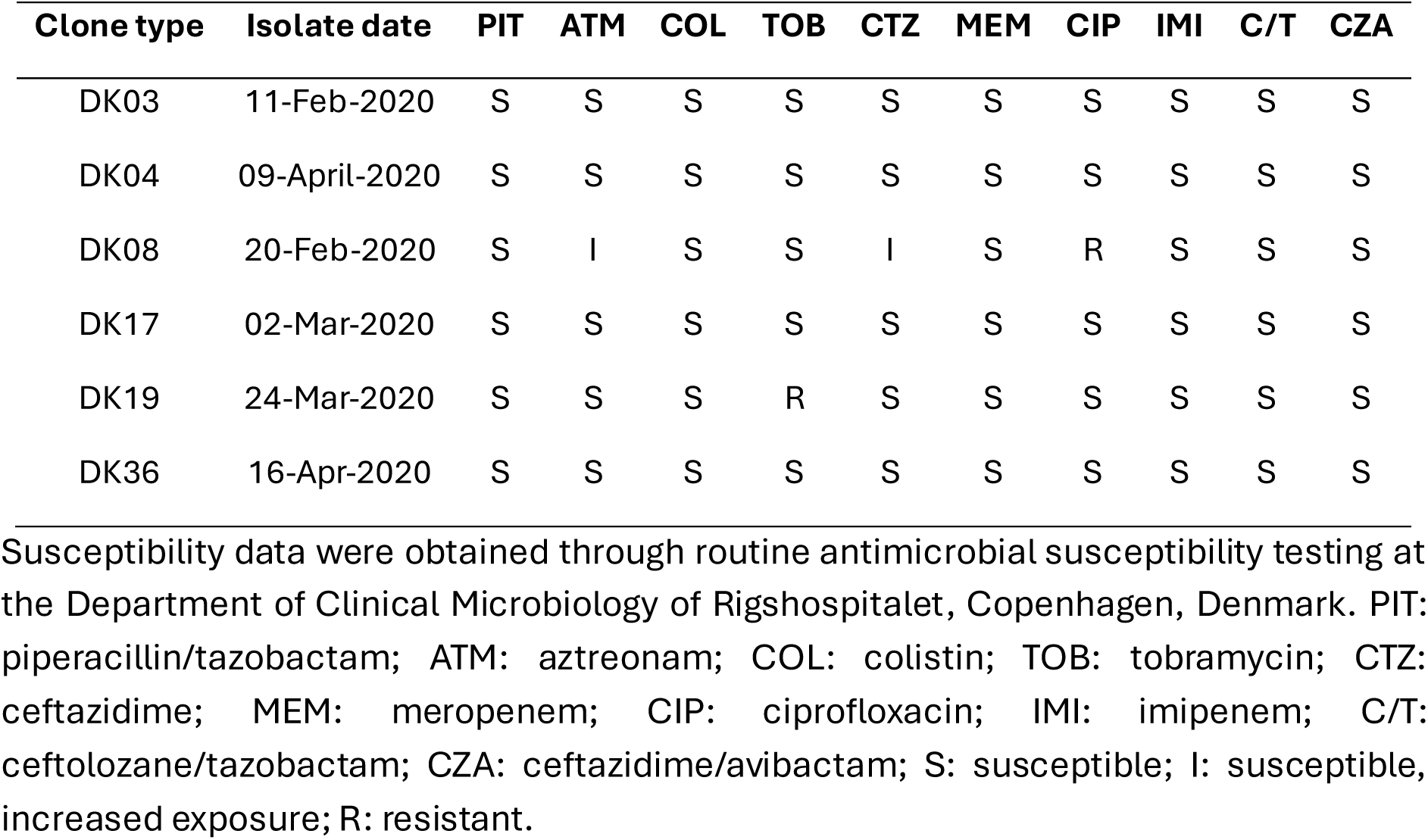
Antimicrobial susceptibility profiles of isolates recovered in 2020 during longitudinal clinical follow-up of the *Pseudomonas aeruginosa* lineages analyzed in this study.

| **Clone type** | **Isolate date** | **PIT** | **ATM** | **COL** | **TOB** | **CTZ** | **MEM** | **CIP** | **IMI** | **C/T** | **CZA** |
|---|---|---|---|---|---|---|---|---|---|---|---|
| DK03 | 11-Feb-2020 | S | S | S | S | S | S | S | S | S | S |
| DK04 | 09-April-2020 | S | S | S | S | S | S | S | S | S | S |
| DK08 | 20-Feb-2020 | S | I | S | S | I | S | R | S | S | S |
| DK17 | 02-Mar-2020 | S | S | S | S | S | S | S | S | S | S |
| DK19 | 24-Mar-2020 | S | S | S | R | S | S | S | S | S | S |
| DK36 | 16-Apr-2020 | S | S | S | S | S | S | S | S | S | S |

Susceptibility data were obtained through routine antimicrobial susceptibility testing at the Department of Clinical Microbiology of Rigshospitalet, Copenhagen, Denmark. PIT: piperacillin/tazobactam; ATM: aztreonam; COL: colistin; TOB: tobramycin; CTZ: ceftazidime; MEM: meropenem; CIP: ciprofloxacin; IMI: imipenem; C/T: ceftolozane/tazobactam; CZA: ceftazidime/avibactam; S: susceptible; I: susceptible, increased exposure; R: resistant.

Whole-genome analysis identified associations rather than causal relationships between mutations and phenotypic outcomes but suggested that these phenotypes arise through multiple genetic routes. Several lineages carried mutations previously associated with classical resistance mechanisms, including alterations in *mexB*, *mpl*, *fusA1* and *nfxB*. However, not all phenotypes could be readily explained by classical resistance mutations alone. Particularly notable was *mexZ*, which was mutated in four of the studied lineages and has previously been reported in up to 40% of clinical *P. aeruginosa* isolates from chronically infected people with CF [2,26]. While *mexZ* mutations are traditionally associated with only modest increases in aminoglycoside MICs [26], previous work demonstrated that disruption of *mexZ* promotes bacterial localization within the airway epithelium, protecting bacteria from antibiotics that poorly penetrate this compartment [8]. The observation that three of the four *mexZ*-mutated lineages displayed enhanced survival in the ALI model without corresponding changes in MIC values is consistent with this mechanism. However, infection-dependent phenotypes were not exclusively associated with increased survival. The DK08 lineage carrying a *nfxB* mutation showed the expected increase in ciprofloxacin resistance but also displayed increased susceptibility to ceftazidime and tobramycin in the ALI model, consistent with previous reports of collateral sensitivity associated with *nfxB*-mediated fluoroquinolone resistance [27–30].

Overall, our results suggest that phenotypes observed under standard laboratory conditions do not uniformly predict those observed in infection-relevant environments, with discordant phenotypes being most apparent among chronically adapted isolates and varying between antibiotics.

## Transparency declaration

### Conffict of interest

The authors declare to have no conflict of interest.

### Funding

This study has been funded by a European Society of Clinical Microbiology and Infectious Diseases (ESCMID) research grant to PL in 2026. The research in the group was funded by a grant from The John and Birthe Meyer Foundation and a Challenge Grant NNF19OC0056411 from The Novo Nordisk Foundation to HKJ.

## Acknowledgements

We thank the Infection Microbiology group for their insightful comments and discussion. The Basal Cell Immortalized Non-Smoker 1.1 (BCi-NS1.1) cell line was a gift from Professor Ronald G. Cristal (Weil Cornell Medical College, New York, USA).

## Contribution

PL designed the study. PL and ECS performed the experimental work. HKJ provided the clinical strains. ECS, RLR, HKJ and SM participated in the design of the study. All authors participated in writing the manuscript and approved the submitted version.

## Notes

### Competing Interest Statement

The authors have declared no competing interest.

