## Supplementary material for "Infection-dependent antibiotic response phenotypes emerge during chronic airway adaptation of *Pseudomonas aeruginosa*"

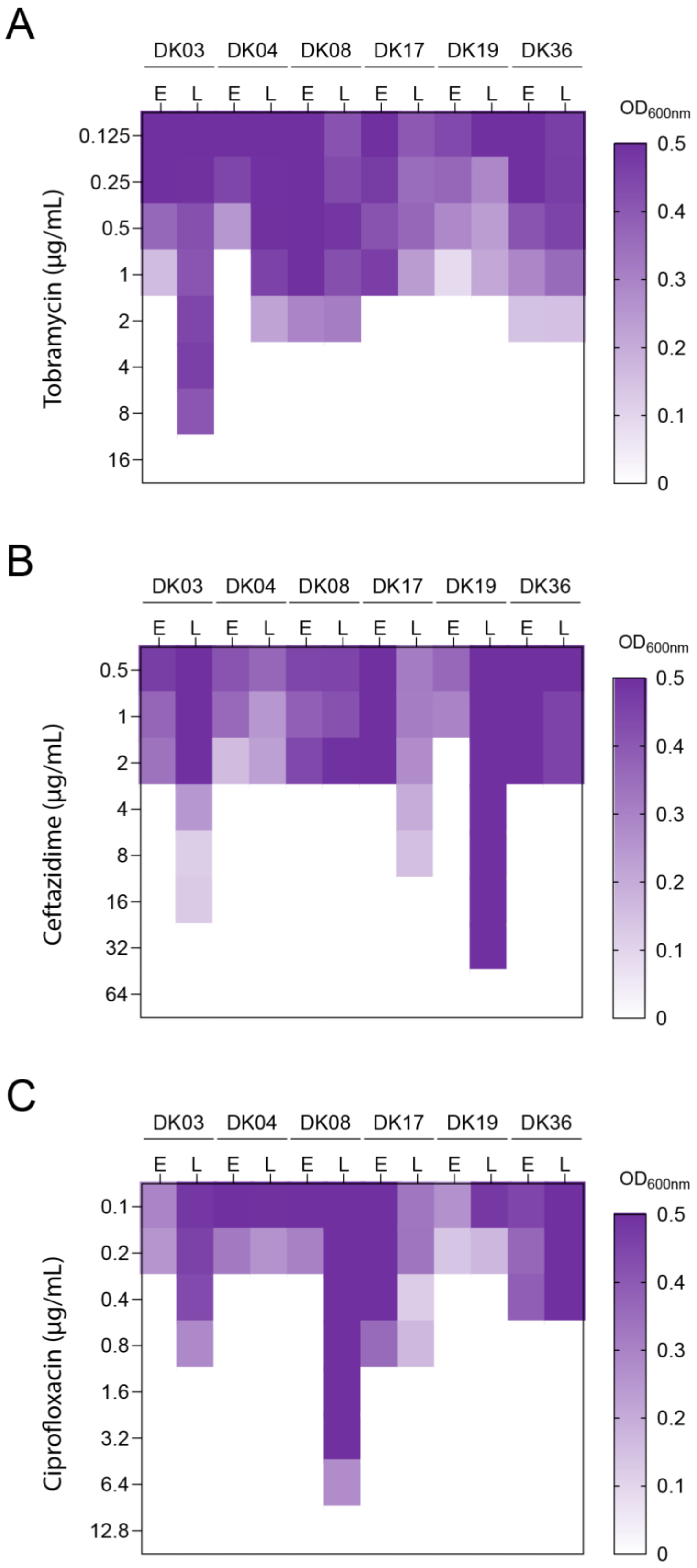


**Supplementary Figure S1. Raw antibiotic susceptibility data for the longitudinal *Pseudomonas aeruginosa* clinical isolates.** Heatmaps show $OD_{600nm}$ measurements obtained after 24 h of broth microdilution assays with increasing concentrations of tobramycin **(A)**, ceftazidime **(B)**, and ciprofloxacin **(C)**. These data were used to determine the MIC values and susceptibility changes during chronic infection reported in Table 1. The reported values are the average of three biological replicates. E: Early strain; L: Late strains.

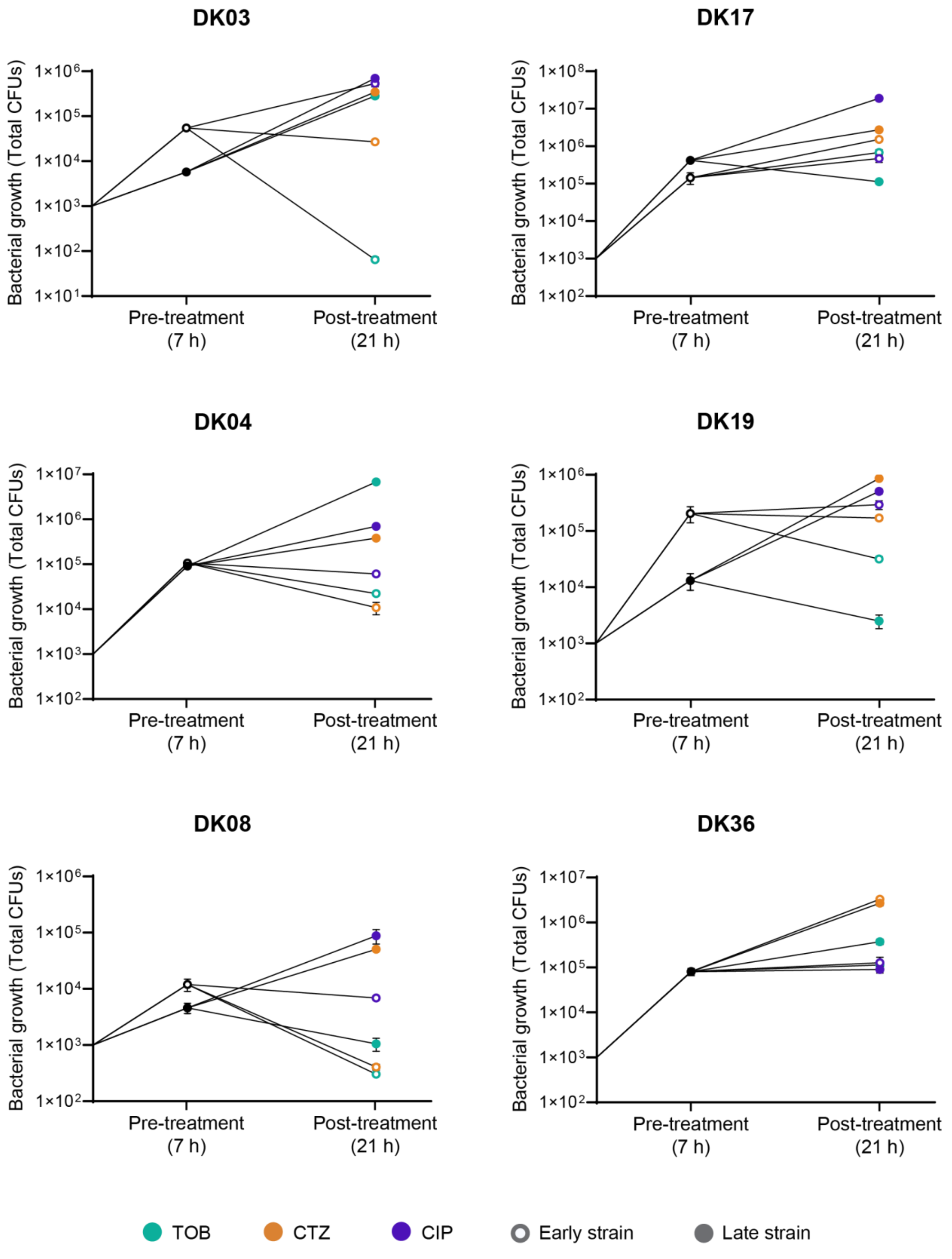


**Supplementary Figure S2. Raw air-liquid interface infection model data used to quantify antibiotic responses.** Total bacterial burden (CFU) was determined immediately before antibiotic treatment (black) and after 14 h of exposure to tobramycin (TOB, green), ceftazidime (CTZ, orange), or ciprofloxacin (CIP, purple) in differentiated airway epithelial cultures. For each clone type, early isolates are represented by open symbols and late isolates by filled symbols. Mean values from three biological replicates are shown, and error bars represent the standard error of the mean (SEM). Lines connect mean CFU values measured before and after antibiotic treatment. These data were used to calculate the fold change in bacterial burden following antibiotic treatment shown in Figure 3A and the corresponding late-to-early response ratios displayed in Figure 3C.

**Supplementary Table 1. Genetic modifications identified in late isolates relative to their corresponding early isolates analyzed in this study.**

| **DK19 (late isolate 163 relative to early isolate 154)** | | | | |
|---|---|---|---|---|
| **Gene name** | **Gene locus** | **Product** | **Mutation type** | **Nucleotide change** |
| *gyrB* | PA0004 | DNA gyrase subunit B | Missense | C1397T |
| *algU* | PA0762 | Sigma factor AlgU | Missense | T419C |
| *cobQ* | PA1277 | Cobyric acid synthase | Silent | C462T |
| - | PA1527 | Conserved hypothetical protein | Silent | T1285C |
| *mexZ* | PA2020 | Multidrug efflux pump regulator | Missense | C530T |
| *xcpQ* | PA3105 | General secretion pathway protein D | Missense | T374C |
| *algG* | PA3545 | Alginate-C5-mannuronan epimerase AlgG | Missense | C1301T |
| *lysP* // - | PA4628 // PA4629 | Intergenic region | Intergenic SNP | A→C |
| *grpE* | PA4762 | Heat shock protein GrpE | Nonsense | G433T |
| - // - | PA0134 // PA0135 | Intergenic region | Intergenic insertion | 153384-153385ins(GCAC) |
| *ilvD* // - | PA0353 // PA0354 | Intergenic region | Intergenic insertion | 398140-398140ins(T) |
| *mexB* | PA0426 | Resistance-Nodulation-Cell Division (RND) multidrug | Deletion | 474932-474944del(GCTCCAGTGCCGA) |

| | | | | |
|---|---|---|---|---|
| | | efflux transporter MexB | | |
| - // - | PA0574.1 // PA0575 | Intergenic region | Intergenic insertion | 630515-630515ins(A) |
| - | PA0640 | Probable bacteriophage protein | Deletion | 694726-694727del(GT) |
| - // - | PA0947 // PA0948 | Intergenic region | Intergenic deletion | 1036123-1036124del(GC) |
| - // - | PA1334 // PA1335 | Intergenic region | Complex indel | +ATT / −CATTCCCCACA |
| *ffs* | PA1530.1 | 4.5S ribosomal RNA | Deletion | 1669081-1669083del(CGT) |
| - | PA1569 | Probable major facilitator superfamily transporter | Insertion | 1710471-1710472ins(CC) |
| - | PA1631 | Probable acyl-CoA dehydrogenase | Insertion | 1774970-1774971ins(AA) |
| - | PA1631 | Probable acyl-CoA dehydrogenase | Insertion | 1774973-1774974ins(G) |
| - | PA2463 | Hypothetical protein | Insertion | 2780431-2780432ins(GGGG) |
| - | PA2824 | Probable sensor/response regulator hybrid | Deletion | 3179447-3179448del(TTT) |
| - | PA2824 | Probable sensor/response regulator hybrid | Deletion | 3179447-3179448del(TTTT) |
| *gdhB* | PA3068 | NAD-dependent | Deletion | 3438199-3438200del(CCCC) |

| | | | | |
|---|---|---|---|---|
| | | glutamate dehydrogenase | | |
| - // - | PA3230 // PA3231 | Intergenic region | Intergenic deletion | 3618697-3618698del(GGCGGA) |
| - // - | PA3468 // PA3469 | Intergenic region | Intergenic insertion | 3882432-3882433ins(GC) |
| - // - | PA3577 // PA3578 | Intergenic region | Intergenic deletion | 4009363-4009364del(C) |
| *mpl* | PA4020 | UDP-N-acetylmuramate:L-alanyl-γ-D-glutamyl-meso-diaminopimelate ligase | Insertion | 4499732-4499733ins(G) |
| *tufA* | PA4265 | Elongation factor Tu | Insertion | 4767834-4767835ins(G) |
| *pykA // -* | PA4329 // PA4330 | Intergenic region | Intergenic insertion | 4858417-4858418ins(CG) |
| - // - | PA4786 // PA4787 | Intergenic region | Intergenic deletion | 5375501-5375502del(CGGG) |
| - // - | PA4873 // PA4874 | Intergenic region | Intergenic insertion | 5471578-5471596ins(CGCCACAGGCGCTATCCGC) |
| - | PA5266 | Conserved hypothetical protein | Deletion | 5930755-5930756del(GCCTGC) |
| *pauB4* | PA5309 | FAD-dependent oxidoreductase | Insertion | 5978719-5978720ins(CCC) |

| **DK36 (late isolate 291 relative to early isolate 277)** | | | | |
|---|---|---|---|---|
| **Gene name** | **Gene locus** | **Product** | **Mutation type** | **Nucleotide change** |
| **-** | PA0272 | Probable transcriptional regulator | Silent | G60A |

| | | | | |
|---|---|---|---|---|
| - | PA3767 | Conserved hypothetical protein | Missense | G74A |
| - // - | PA0588 // PA0589 | Intergenic region | Intergenic insertion | 648705-648706ins(G) |
| - | PA0626 | Hypothetical protein | Insertion | 686213-686213ins(T) |
| - // - | PA0729 // PA0729.1 | Intergenic region | Intergenic insertion | 797719-797720ins(ACA) |
| - // - | PA0729 // PA0729.1 | Intergenic region | Intergenic insertion | 797719-797720ins(GCG) |
| - // - | PA0741 // PA0742 | Intergenic region | Intergenic deletion | 809541-809541del(C) |
| *algU* | PA0762 | Sigma factor AlgU | Insertion | 831429-831430ins(GCCCAGGAA) |
| *mucA* | PA0763 | Anti-sigma factor MucA | Deletion | 832494-832494del(G) |
| - | PA0860 | Probable ATP-binding/permease fusion ABC transporter | Deletion | 938300-938300del(G) |
| *Bcp* | PA1007 // PA1008 | Intergenic region | Intergenic insertion | 1092005-1092006ins(T) |
| - // - | PA1190 // PA1191 | Intergenic region | Intergenic deletion | 1292441-1292442del(C) |
| - // - | PA1194 // PA1195 | Intergenic region | Intergenic deletion | 1296262-1296263del(A) |
| - // - | PA1367 // PA1368 | Intergenic region | Intergenic insertion | 1481083-1481083ins(A) |
| *dnaX* | PA1532 | DNA polymerase subunits gamma and tau | Deletion | 1671363-1671364del(CCGAGC) |
| - // - | PA1606 // PA1607 | Intergenic region | Intergenic deletion | 1749223-1749224del(G) |
| - | PA1796.3 | tRNA-Leu | Insertion | 1947603-1947606ins(CACT) |
| *liuD* | PA2012 | Methylcrotonyl-CoA carboxylase alpha subunit | Deletion | 2201548-2201549del(GGG) |
| *oprI* // - | PA2852.1 // PA2853 | Intergenic region | Intergenic insertion | 3206787-3206787ins(A) |
| - | PA3094 | tRNA-Val | Deletion | 3474708-3474719del(GGAATTCGGCGC) |
| - | PA3297 | Probable ATP-dependent helicase | Deletion | 3692628-3692629del(GGG) |
| - // - | PA3310 // PA3311 | Intergenic region | Intergenic insertion | 3712450-3712451ins(GGGG) |

| | | | | |
|---|---|---|---|---|
| - | PA3346 | Probable two-component response regulator | Insertion | 3757847-3757848ins(CC) |
| - | PA3346 | Probable two-component response regulator | Insertion | 3757847-3757848ins(C) |
| *adk* // *ppc* | PA3686 // PA3687 | Intergenic region | Intergenic insertion | 4127711-4127712ins(C) |
| - // - | PA3690 // PA3691 | Intergenic region | Intergenic insertion | 4133944-4133944ins(T) |
| - | PA4000 | Hypothetical protein | Insertion | 4480458-4480458ins(T) |
| *rplU* // *ispB* | PA4568 // PA4569 | Intergenic region | Intergenic deletion | 5116811-5116812del(C) |
| - | PA4642 | Hypothetical protein | Deletion | 5210907-5210907del(C) |
| - | PA4673 | tRNA-Met | Insertion | 5241832-5241833ins(AA) |
| - // - | PA5132 // PA5133 | Intergenic region | Intergenic insertion | 5780900-5780901ins(T) |
| *pdxY* | PA5516 | Pyridoxamine kinase | Deletion | 6209407-6209408del(TCCACGT) |
| - | PA5532 | Hypothetical protein | Deletion | 6226265-6226266del(CCACCCC) |

| **DK17 (late isolate 499.4 relative to early isolate 484.1)** | | | | |
|---|---|---|---|---|
| **Gene name** | **Gene locus** | **Product** | **Mutation type** | **Nucleotide change** |
| *mexZ* | PA2020 | Multidrug efflux pump regulator | Missense | A29C |
| *pslB* | PA2232 | PslB | Missense | A889G |
| - | PA2480 | Probable two-component sensor | Missense | G970A |
| *clpA* | PA2620 | ATP-binding protease component ClpA | Missense | G433A |
| - // - | PA4104 // PA4105 | Intergenic region | Intergenic SNP | A→C |
| *pchE* | PA4226 | Dihydroaeruginoic acid synthetase | Missense | A1577G |
| - | PA4437 | Hypothetical protein | Missense | A506T |
| - | PA4437 | Hypothetical protein | Missense | G511C |
| - | PA4437 | Hypothetical protein | Missense | A514G |

| | | | | |
|---|---|---|---|---|
| - | PA4437 | Hypothetical protein | Missense | T518A |
| - | PA4656 | Conserved hypothetical protein | Silent | C573T |
| *retS* | PA4856 | RetS (Regulator of Exopolysaccharide and Type III Secretion) | Nonsense | G1531T |
| *ilvD* // - | PA0353 // PA0354 | Intergenic region | Intergenic deletion | 398136-398136del(A) |
| *ilvD* // - | PA0353 // PA0354 | Intergenic region | Intergenic insertion | 398140-398140ins(T) |
| *phhC* | PA0870 | Aromatic amino acid aminotransferase | Deletion | 952120-952121del(GG) |
| *gacS* | PA0928 | Sensor/response regulator hybrid | Deletion | 1013833-1013846del(GCGTTGCAGCTTGC) |
| *pdxB* | PA1375 | Erythronate-4-phosphate dehydrogenase | Insertion | 1493055-1493056ins(GAAT) |
| - // - | PA1377 // PA1378 | Intergenic region | Intergenic insertion | 1495579-1495580ins(CGGAAAAC) |
| *stp1* | PA1670 | Serine/threonine phosphoprotein phosphatase Stp1 | Insertion | 1823215-1823216ins(CTGGGTC) |
| *pscO* | PA1696 | Translocation protein in type III secretion | Deletion | 1845689-1845701del(CGGACGCGCAACA) |
| - // - | PA1841 // PA1842 | Intergenic region | Intergenic insertion | 1999485-1999486ins(CGGTTT) |
| - // - | PA1941 // PA1942 | Intergenic region | Intergenic insertion | 2125968-2125968ins(C) |
| - // *bacA* | PA1958 // PA1959 | Intergenic region | Intergenic deletion | 2142952-2142953del(GGAAAAA) |
| *pcoA* | PA2065 | Copper resistance protein A precursor | Insertion | 2263646-2263660ins(GTCCATGCCGTTCAT) |
| - | PA2160 | Probable glycosyl hydrolase | Insertion | 2379882-2379883ins(AG) |
| - | PA2160 | Probable glycosyl hydrolase | Insertion | 2379888-2379889ins(A) |

| | | | | |
|---|---|---|---|---|
| - | PA2160 | Probable glycosyl hydrolase | Insertion | 2379890-2379891ins(CATTGAGGA) |
| - | PA2160 | Probable glycosyl hydrolase | Insertion | 2379890-2379891ins(GA) |
| - | PA2160 | Probable glycosyl hydrolase | Insertion | 2379891-2379892ins(T) |
| - | PA2330 | Hypothetical protein | Deletion | 2571782-2571783del(G) |
| - | PA2490 | Conserved hypothetical protein | Insertion | 2806290-2806291ins(AACGAAT) |
| *xcpP* | PA3104 | Secretion protein XcpP | Insertion | 3484022-3484023ins(AAAAAAA) |
| *fimV* | PA3115 | Motility protein FimV | Deletion | 3496465-3496474del(GCAGGTCGGT) |
| - // - | PA3230 // PA3231 | Intergenic region | Intergenic insertion | 3618697-3618698ins(GGCGGA) |
| - | PA3297 | Probable ATP-dependent helicase | Insertion | 3693752-3693753ins(CTGGTT) |
| - | PA3297 | Probable ATP-dependent helicase | Insertion | 3693797-3693798ins(TGTAG) |
| - // - | PA3414 // PA3415 | Intergenic region | Intergenic deletion | 3820916-3820917del(CCA) |
| - | PA3453 | Conserved hypothetical protein | Deletion | 3858592-3858593del(C) |
| - // - | PA3496 // PA3497 | Intergenic region | Intergenic insertion | 3914104-3914104ins(T) |
| *moaC* // - | PA3918 // PA3919 | Intergenic region | Intergenic deletion | 4387133-4387134del(GGAC) |
| - // - | PA3927 // PA3928 | Intergenic region | Intergenic deletion | 4403522-4403526del(TCCCG) |
| - | PA3939 | Hypothetical protein | Deletion | 4417078-4417079del(CGAA) |
| - | PA4013 | Conserved hypothetical protein | Deletion | 4493463-4493464del(C) |
| *phzB1* | PA4211 | Probable phenazine biosynthesis protein | Insertion | 4714800-4714801ins(GATC) |
| - | PA4437 | Hypothetical protein | Insertion | 4970098-4970099ins(GGCGCGCCT) |
| - | PA4437 | Hypothetical protein | Insertion | 4970273-4970274ins(TG) |

| | | | | |
|---|---|---|---|---|
| *panC* | PA4730 | Pantoate--beta-alanine ligase | Insertion | 5312996-5312997ins(G) |
| - | PA4798 | Hypothetical protein | Insertion | 5384501-5384502ins(AAT) |
| - // - | PA4802 // PA4802.1 | Intergenic region | Intergenic insertion | 5387754-5387755ins(AGGC) |
| *aceE* | PA5015 | Pyruvate dehydrogenase | Insertion | 5636743-5636744ins(ATCT) |
| *arcC* // - | PA5173 // PA5174 | Intergenic region | Intergenic deletion | 5825788-5825789del(CGCCC) |
| - // - | PA5464 // PA5465 | Intergenic region | Intergenic insertion | 6154750-6154751ins(TC) |

| **DK03 (late isolate 310.1 relative to early isolate 318)** | | | | |
|---|---|---|---|---|
| **Gene name** | **Gene locus** | **Product** | **Mutation type** | **Nucleotide change** |
| *triA* | PA0156 | Resistance-Nodulation-Cell Division (RND) triclosan efflux membrane fusion protein TriA | Missense | A329G |
| - | PA0476 | Probable permease | Missense | T1618C |
| - | PA0547 | Probable transcriptional regulator | Missense | T235C |
| - | PA1153 | Hypothetical protein | Silent | G351A |
| - // - | PA1379 // PA1380 | Intergenic region | Intergenic SNP | T→C |
| *lasR* | PA1430 | Transcriptional regulator LasR | Missense | T649C |
| *phzB2* | PA1900 | Probable phenazine biosynthesis protein | Missense | T412C |
| *mexZ* | PA2020 | Multidrug efflux pump regulator | Missense | T128C |
| - | PA2042 | Probable transporter (membrane subunit) | Silent | T501C |
| *accD* | PA3112 | Acetyl-CoA carboxylase beta subunit | Missense | T52C |

| | | | | |
|---|---|---|---|---|
| *rpsA* | PA3162 | 30S ribosomal protein S1 | Missense | T1129C |
| - | PA3199 | Conserved hypothetical protein | Silent | A414G |
| - | PA3264 | Probable transporter | Silent | C463T |
| - | PA3316 | Probable permease of ABC transporter | Silent | C765T |
| - | PA3754 | Hypothetical protein | Missense | T569C |
| *nagR* | PA3757 | Transcriptional regulator of N-acetylglucosamine catabolism operon | Missense | A83G |
| - | PA3825 | Hypothetical protein | Silent | T1548C |
| *fusA1* | PA4266 | Elongation factor G | Missense | T1786C |
| *inaA* | PA4378 | InaA protein | Missense | T548A |
| - // - | PA0186 // PA0187 | Intergenic region | Intergenic insertion | 213412-213415ins(GGGA) |
| *pcaC* // - | PA0232 // PA0233 | Intergenic region | Intergenic insertion | 262436-262437ins(GGCGGCGC) |
| - | PA0339 | Hypothetical protein | Deletion | 382293-382294del(C) |
| *ilvD* // - | PA0353 // PA0354 | Intergenic region | Intergenic insertion | 398136-398136ins(A) |
| - // - | PA0535 // PA0536 | Intergenic region | Intergenic deletion | 595168-595168del(G) |
| - | PA0818 | Hypothetical protein | Insertion | 895505-895506ins(GTCC) |
| - // - | PA0876 // PA0877 | Intergenic region | Intergenic insertion | 959373-959374ins(ACAGGCTG) |
| - // - | PA0876 // PA0877 | Intergenic region | Intergenic insertion | 959373-959374ins(ACAGGCTGTC) |
| - // - | PA0876 // PA0877 | Intergenic region | Intergenic insertion | 959475-959476ins(GG) |
| - | PA1115 | Hypothetical protein | Deletion | 1209727-1209728del(G) |
| - // - | PA1334 // PA1335 | Intergenic region | Intergenic insertion | 1447060-1447061ins(ATT) |
| *cysC* // - | PA1393 // PA1394 | Intergenic region | Intergenic insertion | 1516315-1516316ins(GA) |
| - | PA1550 | Hypothetical protein | Deletion | 1687798-1687808del(GCACCGCGATG) |

| | | | | |
|---|---|---|---|---|
| - | PA2099 | Probable short-chain dehydrogenase | Deletion | 2311107-2311108del(AC) |
| - // - | PA2427 // PA2428 | Intergenic region | Intergenic deletion | 2723307-2723308del(CA) |
| - | PA2894 | Hypothetical protein | Deletion | 3249252-3249253del(C) |
| *wbpM* // - | PA3141 // PA3142 | Intergenic region | Intergenic insertion | 3526714-3526715ins(TCTG) |
| *zwf* | PA3183 | Glucose-6-phosphate 1-dehydrogenase | Insertion | 3572408-3572409ins(T) |
| - // *cyaB* | PA3216 // PA3217 | Intergenic region | Intergenic insertion | 3605470-3605471ins(G) |
| - | PA3316 | Probable permease of ABC transporter | Deletion | 3719198-3719199del(TCAATAACCT) |
| - // - | PA3468 // PA3469 | Intergenic region | Intergenic insertion | 3882432-3882433ins(GC) |
| - // - | PA3496 // PA3497 | Intergenic region | Intergenic deletion | 3914110-3914111del(GG) |
| *mucP* | PA3649 | MucP | Deletion | 4088197-4088198del(G) |
| - // - | PA4155 // PA4156 | Intergenic region | Intergenic deletion | 4650336-4650337del(CGGCTTCT) |
| - // - | PA4595 // PA4596 | Intergenic region | Intergenic insertion | 5148927-5148928ins(TCATCGTCA) |
| - // - | PA4786 // PA4787 | Intergenic region | Intergenic deletion | 5375504-5375505del(GCA) |

| **DK04 (late isolate 26 relative to early isolate 75)** | | | | |
|---|---|---|---|---|
| **Gene name** | **Gene locus** | **Product** | **Mutation type** | **Nucleotide change** |
| *hemF* | PA0024 | Coproporphyrinogen III oxidase, aerobic | Silent | C744T |
| *mexZ* | PA2020 | Multidrug efflux pump regulator | Missense | T488C |
| - // - | PA2497 // PA2498 | Intergenic region | Intergenic SNP | G→A |
| *phzB1* | PA4211 | Probable phenazine biosynthesis protein | Silent | T486C |
| *pnp* | PA4740 | Polyribonucleotide nucleotidyltransferase | Silent | T1977C |

| | | | | |
|---|---|---|---|---|
| - | PA4880 | Probable bacterioferritin | Missense | A443C |
| - | PA5335 | Conserved hypothetical protein | Missense | G110C |
| *algB* | PA5483 | Two-component response regulator AlgB | Silent | G549A |
| *gyrB* // *lptA* | PA0004 // PA0005 | Intergenic region | Intergenic insertion | 6772-6782ins(CGGGCGCCAGG) |
| - | PA0259 | Hypothetical protein | Deletion | 289663-289664del(G) |
| *ilvD* // - | PA0353 // PA0354 | Intergenic region | Intergenic insertion | 398136-398136ins(A) |
| *ilvD* // - | PA0353 // PA0354 | Intergenic region | Intergenic deletion | 398140-398140del(T) |
| - // - | PA0560 // PA0561 | Intergenic region | Intergenic deletion | 618152-618153del(C) |
| *apaH* | PA0590 | Bis(5'-nucleosyl)-tetraphosphatase | Deletion | 649872-649873del(CC) |
| - | PA1239 | Hypothetical protein | Deletion | 1341375-1341376del(GAGTGGCCGATC) |
| - // - | PA2029 // PA2030 | Intergenic region | Intergenic deletion | 2220598-2220599del(GGGAGGGTGG) |
| - // - | PA2029 // PA2030 | Intergenic region | Intergenic deletion | 2220598-2220599del(GGGTGG) |
| - | PA2057 | Hypothetical protein | Deletion | 2251287-2251288del(C) |
| - | PA3290 | Hypothetical protein | Insertion | 3682825-3682826ins(GGTTTCAGGCGT) |
| - // - | PA3457 // PA3458 | Intergenic region | Intergenic deletion | 3865155-3865156del(CGCCG) |
| - // - | PA3496 // PA3497 | Intergenic region | Intergenic insertion | 3914104-3914104ins(T) |
| - // - | PA3577 // PA3578 | Intergenic region | Intergenic deletion | 4009458-4009459del(CT) |
| - | PA4621 | Probable oxidoreductase | Insertion | 5181492-5181493ins(G) |
| - // - | PA4786 // PA4787 | Intergenic region | Intergenic deletion | 5375501-5375502del(CGGG) |
| *pilM* | PA5044 | Type IV fimbrial biogenesis protein PilM | Deletion | 5680176-5680177del(A) |
| *pdxY* | PA5516 | Pyridoxamine kinase | Insertion | 6209407-6209408ins(TCCACGT) |

| DK08 (late isolate 511.1 relative to early isolate 512) | | | | |
|---|---|---|---|---|
| **Gene name** | **Gene locus** | **Product** | **Mutation type** | **Nucleotide change** |
| *fmt* | PA0018 | Methionyl-tRNA formyltransferase | Missense | C835G |
| *fmt* | PA0018 | Methionyl-tRNA formyltransferase | Missense | C841G |
| *fmt* | PA0018 | Methionyl-tRNA formyltransferase | Missense | C844T |
| *fmt* | PA0018 | Methionyl-tRNA formyltransferase | Missense | C847A |
| *fmt* | PA0018 | Methionyl-tRNA formyltransferase | Missense | T853C |
| *fmt* | PA0018 | Methionyl-tRNA formyltransferase | Deletion | 20930-20931del(C) |
| *vgrG1a* | PA0091 | T6SS spike protein VgrG1a | Silent | C555G |
| *vgrG1a* | PA0091 | T6SS spike protein VgrG1a | Missense | G1415T |
| *vgrG1a* | PA0091 | T6SS spike protein VgrG1a | Missense | T1416G |
| *eagT6* | PA0094 | T6SS effector | Missense | G340T |
| *eagT6* | PA0094 | T6SS effector | Missense | G349T |
| *eagT6* | PA0094 | T6SS effector | Missense | G358T |
| *vgrG1b* | PA0095 | T6SS spike protein VgrG1b | Silent | G1242C |
| *vgrG1b* | PA0095 | T6SS spike protein VgrG1b | Silent | C1254G |
| - // - | PA0118 // PA0119 | Intergenic region | Intergenic SNP | C→G |
| *mdcH* | PA0214 | Malonate decarboxylase subunit epsilon | Silent | C411T |
| *mdcH* | PA0214 | Malonate decarboxylase subunit epsilon | Missense | C406A |
| *mdcH* | PA0214 | Malonate decarboxylase subunit epsilon | Silent | T402C |

| | | | | |
|---|---|---|---|---|
| *mdcH* | PA0214 | Malonate decarboxylase subunit epsilon | Missense | C397T |
| *hudA* | PA0254 | Pyrrole-2-carboxylic acid decarboxylase | Missense | C112A |
| *hudA* // - | PA0254 // PA0255 | Intergenic region | Intergenic SNP | T→C |
| - | PA0799 | ATP-dependent helicase | Missense | G1786A |
| - | PA0827 | Hypothetical protein | Missense | C19A |
| *aruG* | PA0897 | Arginine N-succinyltransferase | Missense | C916G |
| *ruvC* | PA0965 | Crossover junction endodeoxyribonuclease | Missense | C473T |
| *ruvC* | PA0965 | Crossover junction endodeoxyribonuclease | Missense | G480T |
| *ruvC* | PA0965 | Crossover junction endodeoxyribonuclease | Missense | G488T |
| *ruvC* | PA0965 | Crossover junction endodeoxyribonuclease | Missense | G493C |
| *ruvC* | PA0965 | Crossover junction endodeoxyribonuclease | Missense | G495T |
| *ruvC* | PA0965 | Crossover junction endodeoxyribonuclease | Missense | C499G |
| *ruvC* | PA0965 | Crossover junction endodeoxyribonuclease | Missense | G503T |
| *ruvC* | PA0965 | Crossover junction endodeoxyribonuclease | Missense | G507C |
| *ruvC* | PA0965 | Crossover junction endodeoxyribonuclease | Insertion | 1051633-1051634ins(T) |

| | | | | |
|---|---|---|---|---|
| - // *cobV* | PA1280 // PA1281 | Intergenic region | Intergenic SNP | C→G |
| *cobD* | PA1275 | Threonine-phosphate decarboxylase | Missense | C691G |
| *cobD* | PA1275 | Threonine-phosphate decarboxylase | Missense | A695C |
| *cobD* | PA1275 | Threonine-phosphate decarboxylase | Missense | C709A |
| *cobD* | PA1275 | Threonine-phosphate decarboxylase | Missense | C748G |
| *hxuR* | PA1301 | Putative transmembrane sensor | Silent | C855T |
| *hxuR* | PA1301 | Putative transmembrane sensor | Missense | G862A |
| *hxuR* | PA1301 | Putative transmembrane sensor | Missense | G865C |
| *fliO* | PA1445 | Flagellar protein | Missense | G280T |
| *ccmB* // *ccmC* | PA1476 // PA1477 | Intergenic region | Intergenic SNP | C→G |
| *ccmB* // *ccmC* | PA1476 // PA1477 | Intergenic region | Intergenic SNP | G→C |
| - | PA1558 | Hypothetical protein | Silent | G555A |
| *ccoN* | PA1557 | Cytochrome-c oxidase, cbb3-type subunit I | Silent | T906C |
| *ccoN* | PA1557 | Cytochrome-c oxidase, cbb3-type subunit I | Silent | C921A |
| - | PA1591 | Hypothetical protein | Silent | C381T |
| *pscU* | PA1690 | T3SS export apparatus subunit | Missense | T673C |
| *pscO* | PA1696 | Translocation protein in T3SS | Missense | G433T |
| *pscO* | PA1696 | Translocation protein in T3SS | Missense | A436G |
| *pscO* | PA1696 | Translocation protein in T3SS | Missense | A439G |
| *pscO* | PA1696 | Translocation protein in T3SS | Missense | C442T |

| | | | | |
|---|---|---|---|---|
| *pscO* | PA1696 | Translocation protein in T3SS | Missense | C445A |
| *exsA* | PA1713 | T3SS regulon transcriptional activator ExsA | Insertion | 1857422-1857423ins(GCCAGGC) |
| - | PA1865 | Fanconi-associated nuclease 1-like protein | Missense | G1546A |
| *ercS'* | PA1976 | Hybrid sensor histidine kinase/response regulator | Missense | C2491A |
| *ercS'* | PA1976 | Hybrid sensor histidine kinase/response regulator | Insertion | 2162858-2162859ins(G) |
| *ercS'* | PA1976 | Hybrid sensor histidine kinase/response regulator | Missense | C2498T |
| *ercS'* | PA1976 | Hybrid sensor histidine kinase/response regulator | Deletion | 2162865-2162865del(G) |
| *ercS'* | PA1976 | Hybrid sensor histidine kinase/response regulator | Missense | C2509T |
| *ercS'* | PA1976 | Hybrid sensor histidine kinase/response regulator | Insertion | 2162880-2162881ins(C) |
| *ercS'* | PA1976 | Hybrid sensor histidine kinase/response regulator | Deletion | 2162887-2162887del(T) |
| *ercS'* | PA1976 | Hybrid sensor histidine kinase/response regulator | Silent | C2526T |
| *ercS'* | PA1976 | Hybrid sensor histidine kinase/response regulator | Missense | A2536C |
| *hbcR* | PA2005 | Sigma-54-dependent transcriptional regulator | Silent | G1A |
| *hbcR* | PA2005 | Sigma-54-dependent | Missense | A8C |

| | | | | |
|---|---|---|---|---|
| | | transcriptional regulator | | |
| *hbcR* | PA2005 | Sigma-54-dependent transcriptional regulator | Missense | C14G |
| *hbcR* | PA2005 | Sigma-54-dependent transcriptional regulator | Missense | A16G |
| *hbcR* // - | PA2005 // PA2006 | Intergenic region | Intergenic SNP | G→C |
| *hbcR* // - | PA2005 // PA2006 | Intergenic region | Intergenic SNP | T→C |
| *hbcR* // - | PA2005 // PA2006 | Intergenic region | Intergenic SNP | G→A |
| *hbcR* // - | PA2005 // PA2006 | Intergenic region | Intergenic SNP | G→C |
| *kynB* | PA2081 | Kynurenine formamidase | Missense | T451G |
| - // *pslA* | PA2230 // PA2231 | Intergenic region | Intergenic SNP | T→C |
| - | PA2286 | Hypothetical protein | Missense | C529A |
| - | PA2422 | Hypothetical protein | Silent | T264C |
| - | PA2452 | Hypothetical protein | Silent | G543C |
| - | PA2452 | Hypothetical protein | Missense | C547A |
| - | PA2452 | Hypothetical protein | Missense | C553G |
| - | PA2452 | Hypothetical protein | Silent | T561C |
| - | PA2452 | Hypothetical protein | Missense | G563T |
| - | PA2452 | Hypothetical protein | Missense | C563G |
| - | PA2452 | Hypothetical protein | Missense | C568A |
| - | PA2452 | Hypothetical protein | Missense | G572C |
| - | PA2452 | Hypothetical protein | Missense | T577G |
| - | PA2452 | Hypothetical protein | Missense | A578C |
| *muxA* | PA2528 | Multidrug efflux RND transporter periplasmic adaptor | Missense | G98T |

| | | | | |
|---|---|---|---|---|
| - | PA2679 | Hypothetical protein | Insertion | 3026793-3026794Ins(G) |
| - | PA2820 | Hypothetical protein | Silent | C198T |
| - | PA2835 | Probable major facilitator superfamily (MFS) transporter | Missense | G1159C |
| *atuR* | PA2885 | Putative repressor of atu genes | Deletion | 3237312-3237313del(T) |
| *cobN* | PA2944 | Cobaltochelatase subunit | Missense | C2422A |
| *acpP // fabG* | PA2966 // PA2967 | Intergenic region | Intergenic SNP | T→G |
| *fimV // -* | PA3115 // PA3116 | Intergenic region | Intergenic SNP | A→G |
| *gyrA* | PA3168 | DNA gyrase subunit A | Missense | A247G |
| *gyrA* | PA3168 | DNA gyrase subunit A | Missense | A751G |
| *ykgJ* | PA3209 | YkgJ family cysteine cluster protein | Silent | G12A |
| - | PA3312 | Probable 3-hydroxyisobutyrate dehydrogenase | Silent | C804G |
| - | PA3312 | Probable 3-hydroxyisobutyrate dehydrogenase | Missense | A823C |
| - | PA3312 | Probable 3-hydroxyisobutyrate dehydrogenase | Silent | C834G |
| - | PA3312 | Probable 3-hydroxyisobutyrate dehydrogenase | Missense | G836T |
| - | PA3421 | Probable ferredoxin | Missense | G2115T |
| - | PA3421 | Probable ferredoxin | Missense | C2116G |
| - | PA3421 | Probable ferredoxin | Missense | T2119C |
| - | PA3421 | Probable ferredoxin | Missense | G2120A |
| *ssuB* | PA3442 | Aliphatic sulfonates | Missense | A580C |

| | | | | |
|---|---|---|---|---|
| | | import ATP-binding protein | | |
| - // - | PA3577 // PA3578 | Intergenic | Intergenic SNP | G→C |
| *adhC* | PA3629 | Class III alcohol dehydrogenase | Missense | A166C |
| *yejK* | PA3849 | Nucleoid-associated protein | Missense | C85T |
| - | PA4039 | Hypothetical protein | Missense | G643T |
| *rpoN* | PA4462 | RNA polymerase factor sigma-54 | Deletion | 4993119-4993138del(GCCCAGCGTCGAAAGCCTCG) |
| *magD* | PA4489 | Alpha-2-macroglobulin | Missense | G3979C |
| *mexC // nfxB* | PA4599 // PA4600 | Intergenic region | Intergenic SNP | G→A |
| *nfxB* | PA4600 | Efflux pump transcriptional repressor | Deletion | 5155884-5155896del(CGCACGGCGAAGG) |
| *cdrA* | PA4625 | Cyclic diguanylate-regulated TPS partner A | Missense | G717A |
| - | PA4689 | Hypothetical protein | Silent | G1020A |
| - | PA4724 | Probable aminoacyl-transfer RNA synthetase (class I) | Missense | G672C |
| - | PA4724 | Probable aminoacyl-transfer RNA synthetase (class I) | Missense | C695A |
| *desB* | PA4888 | Fatty acid desaturase | Silent | C1053G |
| *desB* | PA4888 | Fatty acid desaturase | Silent | C1056G |
| - | PA5234 | Probable oxidoreductase | Missense | G490A |
| - | PA5252 | ATP-binding component of ABC transporter | Missense | G1275C |
| *dgcH* | PA5487 | Diguanylate cyclase | Silent | A354C |
